# SIMILE enables alignment of fragmentation mass spectra with statistical significance

**DOI:** 10.1101/2021.02.24.432767

**Authors:** Daniel G.C. Treen, Trent R. Northen, Benjamin P. Bowen

**Affiliations:** Environmental Genomics and Systems Biology Division & The Joint Genome Institute, Lawrence Berkeley National Laboratory, One Cyclotron Road, Berkeley, CA, 94720

## Abstract

Interrelating compounds according to their aligned fragmentation spectra is central to tandem mass spectrometry-based metabolomics. However, current alignment algorithms do not provide statistical significance and compounds that have multiple delocalized structural differences often fail to have their fragment ions aligned. Significant Interrelation of MS/MS Ions via Laplacian Embedding (SIMILE) is a new tool inspired by protein sequence alignment for aligning fragmentation spectra with statistical significance and allowance for multiple chemical differences. We found SIMILE yields 550% more pairs of structurally similar compounds than commonly used cosine-based scoring algorithms, and anticipate SIMILE will fill an important role by also providing p-values for fragmentation spectra alignments to explore structural relationships between compounds.

## Introduction

Tandem mass spectrometry is widely used in metabolomics experiments to help make chemical assignments. Compound identification typically requires determining if an experimental fragmentation spectrum matches an authentic standard. This is done by aligning fragment ions that share the same mass-to-charge ratio (*m/z*) and calculating the cosine similarity of their intensities^1^. Recently, more generalized alignment approaches have been developed that aim to yield scores that are a better proxy for compound similarity rather than identity. For instance, GNPS molecular networking and NIST Hybrid Search both implement an alignment approach that is sensitive to compounds that differ by a single/localized structural difference(s)^2,3^. This is done by allowing alignment of fragment ions that either share the same *m/z* or the same neutral-loss relative to their intact precursor ion. Machine learning approaches such as SIRIUS, CANOPUS, MS2LDA, and Spec2Vec also incorporate precursor ion neutral-losses as a feature in their implementations^4–7^. While these approaches have shown great success and there are methods for estimating statistical significance for compound identification, to our knowledge no method for calculating the significance of fragmentation spectra alignments has been described^8^.

By contrast, protein sequence alignment methods based on the Needleman-Wunsch algorithm such as BLAST yield alignments with statistical significance that are robust to multiple substitutions, insertions, and deletions^9,10^. These methods are fundamentally different from fragmentation spectra based cosine similarity in that they rely on substitution matrices describing the log-odds of amino acids sharing common ancestry relative to random chance such as the PAM and BLOSUM matrices^11,12^. Unlike protein substitution matrices which are generally of size 20 by 20 (amino acids), a global substitution matrix for fragment ions would be infinite due to the infinite number of possible *m/z* values. In addition, *m/z* values are only partially tied to chemical structure due to the one-to-many correspondence between *m/z* values and chemical structures. However, if restricted to a single pair of fragmentation spectra, a spectral graph-theoretic framework parameterized by their all-by-all *m/z* difference counts can generate finite, context sensitive, and mathematically consistent fragment ion substitution matrices based on expected hitting-times^13^.

Here, we introduce Significant Interrelation of MS/MS Ions via Laplacian Embedding (SIMILE), an approach that leverages methods used for protein sequence alignment to enable robust pairwise alignment of fragmentation spectra with p-value estimation (Fig. 1). Rather than requiring identical *m/z* values or precursor ion neutral-losses for alignment of fragmentation spectra, SIMILE uses all *m/z* differences among a pair of fragmentation spectra to generate a pair specific fragment ion substitution matrix. This matrix is then used as the input to a dynamic programming alignment algorithm for alignment and scoring. The significance of an alignment is calculated via a Monte Carlo permutation test with alignment score as the test statistic under the null hypothesis that *m/z* values are exchangeable between *m/z* ordered fragmentation spectra only if they yield hypothetical fragmentation spectra which are also *m/z* ordered.

**Fig. 1.**
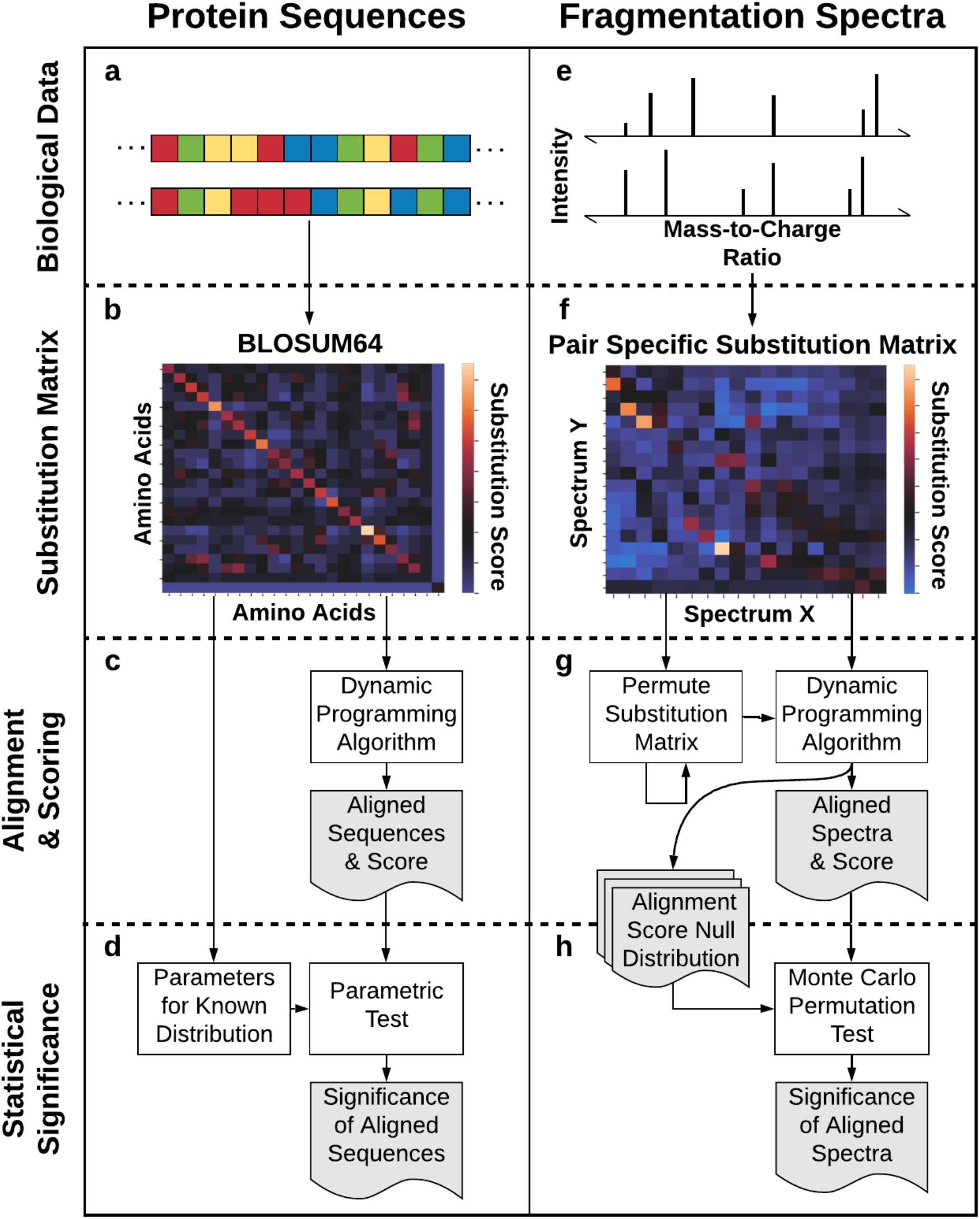
Analogous to how protein sequences undergo alignment, SIMILE aligns fragmentation spectra with allowances for substitutions and gaps. **a-c)** Pairs of protein sequences are globally aligned by using a dynamic programming algorithm such as Needleman-Wunsch with an evolutionary model-based substitution matrix and gap penalty. **d)** Significance of the alignment is calculated by assuming alignment scores follow a distribution parameterized by choice of substitution matrix. **e-g)** Likewise, pairs of fragmentation spectra are aligned by a dynamic programming algorithm with a pair specific substitution matrix and a gap penalty of zero. **g-h)** The alignment score null distribution used to calculate the significance of observed alignments stems from permuting pair specific substitution matrix interrelating fragmentation spectra X and Y randomly many times with restrictions according to the null hypothesis.

## Methods

### Overview

The SIMILE algorithm calculates spectral similarity by aligning substitutable fragments; identification of the significance of their alignment; and scoring the degree to which two spectra are correlated. The underlying mathematical framework for these calculations is described below. Python code for each step is available in the GitHub repository (https://github.com/biorack/simile). Figure 2 illustrates the SIMILE algorithm with two hypothetical molecules.

**Fig. 2.**
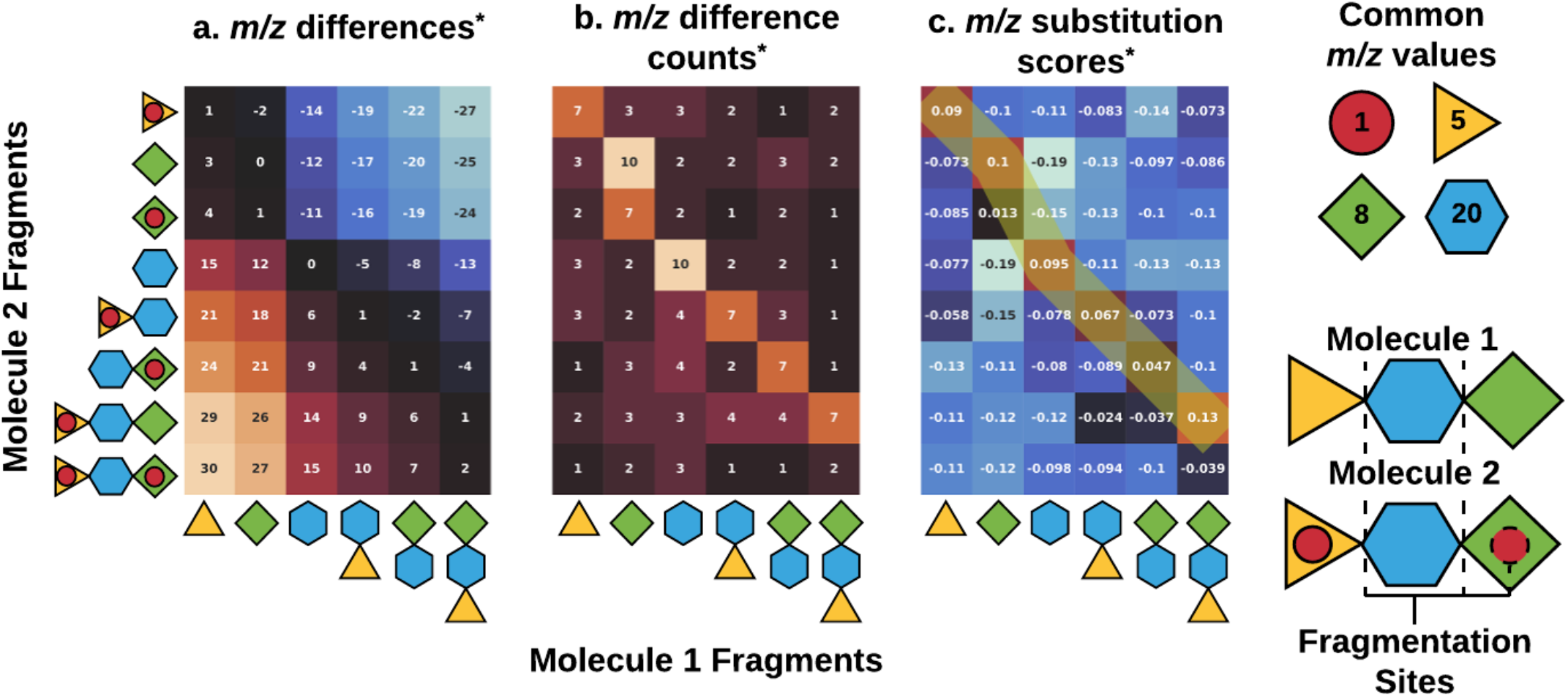
For two hypothetical molecules, this cartoon illustrates the underlying steps of the SIMILE algorithm in calculating pairwise *m/z* substitution matrices. For this example, two molecules “1” and “2” are shown. Each molecule is made from three blocks and there are two modifiers to molecule 2. **a)** All pairwise *m/z* differences between fragments of molecule 1 and molecule 2 are stored in a matrix. **b)** All entries with the same *m/z* difference in the top two quadrants are replaced by the number of entries in which that *m/z* difference occurred. The same process is repeated for the bottom two quadrants. **c)** Calculating the pseudo-inverse of the directed graph laplacian with this matrix yields substitution scores for each pair of *m/z* values. The quadrants interrelating molecule 1 and 2 can then be fed into a dynamic programming algorithm to yield aligned fragment ions between molecule 1 and 2. *\*For illustrative purposes, only the quadrant corresponding to molecule 1 vs. molecule 2 of the full matrices is shown. The other quadrants corresponding to molecule 1 vs. molecule 1, molecule 2 vs. molecule 1, and molecule 2 vs. molecule 2 are still used for calculations.*

### Substitution matrix

Construction of a pair-specific *m/z* substitution matrix takes the concatenation of two ordered *m/z* lists from fragmentation spectra X of length *m* and Y of length *n* as input. The pairwise (outer) difference of the concatenated *m/z* list is stored in an *m+n* by *m+n m/z* difference matrix, **D**. (Fig 2a). The *m/z* difference matrix **D** is delineated by four quadrants: X minus X (e.g. all of the intra-spectral differences in X) in the top-left m by m, Y minus Y (e.g. all of the intra-spectral differences in Y) in the bottom-right n by n, X minus Y in the top-right m by n, and Y minus X (e.g. all of the inter-spectral differences between X and Y) in the bottom-left n by m. In Fig 2, only the top-right quadrant is shown to illustrate *m/z* differences, frequencies of specific differences and ion substitution scores comparing the two hypothetical molecules, but all four quadrants are used in the calculation. For each element **D**_ij_ of the top m by m+n quadrants of **D**, the *m/z* differences in the block that are within a 0.01 *m/z* diameter window of **D**_ij_ are counted and element-wise assigned to the *m/z* difference frequency matrix, **C** (Fig 2b). Repeat the procedure to fill the bottom n by m+n quadrants of **C** using the bottom m by m+n block of **D.**

To calculate a substitution matrix from the *m/z* frequency matrix, we used the approach described by Li et al^13^. Dividing each row of *m/z* difference count matrix **C** by the sum of the row yields to *m/z* transition matrix **T**. Each element **T**_ij_ describes the probability of the *i*^th^ *m/z* from the concatenated *m/z* list transitioning to the *j*^th^ *m/z*. As such, the transition matrix defines a Markov chain representation of the *m/z* values from fragmentation spectra X and Y. Let **I** be the m+n by m+n identity matrix, defined as having ones along the diagonal and zeros elsewhere. Let **p** be the stationary probability distribution of the *m/z* transition matrix **T**, defined by the property that **pT** = **p**. The stationary probability distribution **p** is calculated by dividing the principal eigenvector of **T** by its sum. The directed graph laplacian **L** of **C** is defined as **p^1/2^(I-T)p^-1/2^**. Finally, the *m/z* substitution matrix **S** is calculated by taking the Moore-Penrose pseudoinverse of **L. S^XX^** and **S^YY^** are defined as the top-left m by m quadrant and the bottom-right n by n quadrant of **S** respectively. Likewise, **S^XY^** and **S^YX^** are defined as the top-right m by n quadrant and the bottom-left n by m quadrant of **S** respectively.

### Alignment and alignment score

The alignment and alignment score are calculated using the dynamic programming approach described by Gotoh for sequence alignments, but with a fixed gap-penalty of zero^14^. It is defined by the recurrence relation

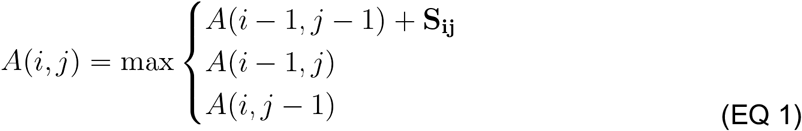

with initial conditions *A*(i,0) = 0 for all i and *A*(0,j) = 0 for all j. The terminal value of *A* is the alignment score and tracing back the elements of **S** which contribute to this score yields the alignment. Because **S^XY^** and **S^YX^** are not necessarily symmetric, we calculate the average alignment score and average alignment length using both quadrants.

### Alignment significance

The significance of the alignment is calculated via a permutation test with the average alignment score used as test statistic under the null hypothesis that *m/z* values are exchangeable between *m/z* ordered fragmentation spectra X and Y only if they generate fragmentation spectra X’ and Y’ which are also *m/z* ordered. This null hypothesis corresponds to the assumption that exchanging inter-spectral substitution scores in **S^XY^** and **S^YX^** with intra-spectral substitution scores in **S^XX^** and **S^YY^** improves alignment scores so long as *m/z* order is preserved in the resulting **S^X^’^Y^’** and **S^Y^’^X^’**.

Rather than compute every valid permutation of **S** to calculate the p-value, we use Monte Carlo testing with early stopping to asymptotically approach the true p-value with bounded and known error^15^. In practice, this is done by first generating a random permutation of indices that index the concatenated *m/z* list such that the first *n* entries are *m/z* ordered and the last *m* entries are *m/z* ordered which implicitly generates hypothetical fragmentation spectra X’ and Y’. This permutation of indices is then used to permute the rows and columns of **S** symmetrically. The average alignment score is calculated using **S^X^’^Y^’** and **S^Y^’^X^’** with the alignment algorithm described above. The p-value is then the probability that a random alignment score from the empirical distribution is greater than the observed alignment score or one out of the number of iterations, whichever is greater (Fig 1h).

### Validation and development dataset

The algorithm was developed and validated using fragmentation spectra from the commercially available electrospray ionization spectra available from the National Institute of Standards and Technology (NIST 2020) library. These spectra are acquired on a variety of instruments under a variety of conditions and were filtered to include only, [M-H]^-^, and collision energy within 5 eV of 40 eV.

Each compound was assigned to a chemical class using the ClassyFire web service^16^. For each unique inchi key, a JSON file containing the chemical class was retrieved from the web service by the following URL http://classyfire.wishartlab.com/entities/ik.json where ik is the inchi key for the compound. While this JSON file provides predicted compound kingdom, superclass, class, and subclass only the class was retained for this analysis.

### Modified cosine spectral similarity

The MatchMS python package version 0.6.2 was used to calculate the modified cosine spectral similarity score and determine the number of matching ions^17^. Each spectrum was square-root intensity scaled and further normalized using the MatchMS function “normalize_intensities”. Pairs of spectra were evaluated with the modified_cosine.pair() function.

### Maximum Common Substructure (MCS)

The similarity metric used in this paper is the Jaccard similarity coefficient, *s*, of the overlapping bonds given by *s* = *N* / ((*N_A_+N_B_*) − *N)* where *N* is the number of overlapping bonds between the pair of molecules *A* and *B*. *N_A_* and *N_B_* are the number of bonds in each of the molecules A and B, respectively. The alignment of the molecules is done using the RDKIT python package with a timeout of 10 seconds and ringMatchesRingOnly set to False^18^. A small number of compound similarities timed out and were discarded from further analysis. Here we will refer to the MCS Jaccard similarity of bonds as MCS.

## Results

### SIMILE p-value filtering

We used fragmentation spectra from NIST-20’s Small Molecule High Resolution Accurate Mass MS/MS Library to determine alignment score cut-offs and p-values that minimize false discovery rates. The fragmentation spectra were filtered by unique InChIKey closest to 40 eV within ± 5 eV to obtain a spectrum for each of 7,356 molecules as negative mode [M-H]- adducts. We did not perform any filtering of ions within spectra. For each dataset, we performed an all-vs-all calculation for modified cosine, SIMILE, and maximum common substructure (MCS) Jaccard similarity coefficient for compound similarity. The average MCS similarity plus one standard deviation was found to be 0.4. Thus, we consider similar structures to be associations where the MCS is greater or equal to than 0.4 and dissimilar structures to be where the MCS is less than 0.4.

From the all-by-all comparisons, we evenly sampled 200,000 comparisons per compound class; where at least one compound in each comparison is in a specific compound class. There were 36 unique compound classes that had at least 200,000 comparisons. Compound classes with less than 200,000 comparisons were not considered for further analysis. These 7.2M comparisons had approximately 1.1M pairs of molecules that were structurally similar (MCS≥0.4). This approach aids to remove bias in the dataset where a small number of compound classes are overrepresented. As shown in Fig 3, the SIMILE p-value and alignment score complement each other to increase the quality of returned associations, where an alignment score cutoff higher than 0.5 and p-value lower than 0.02 gives approximately 20k structurally similar pairs. With these cutoffs, 75% of the pairs identified were actually structurally similar. Selecting alignment score and p-value cutoffs that enable greater specificity comes at the cost of lower sensitivity.

**Fig. 3.**
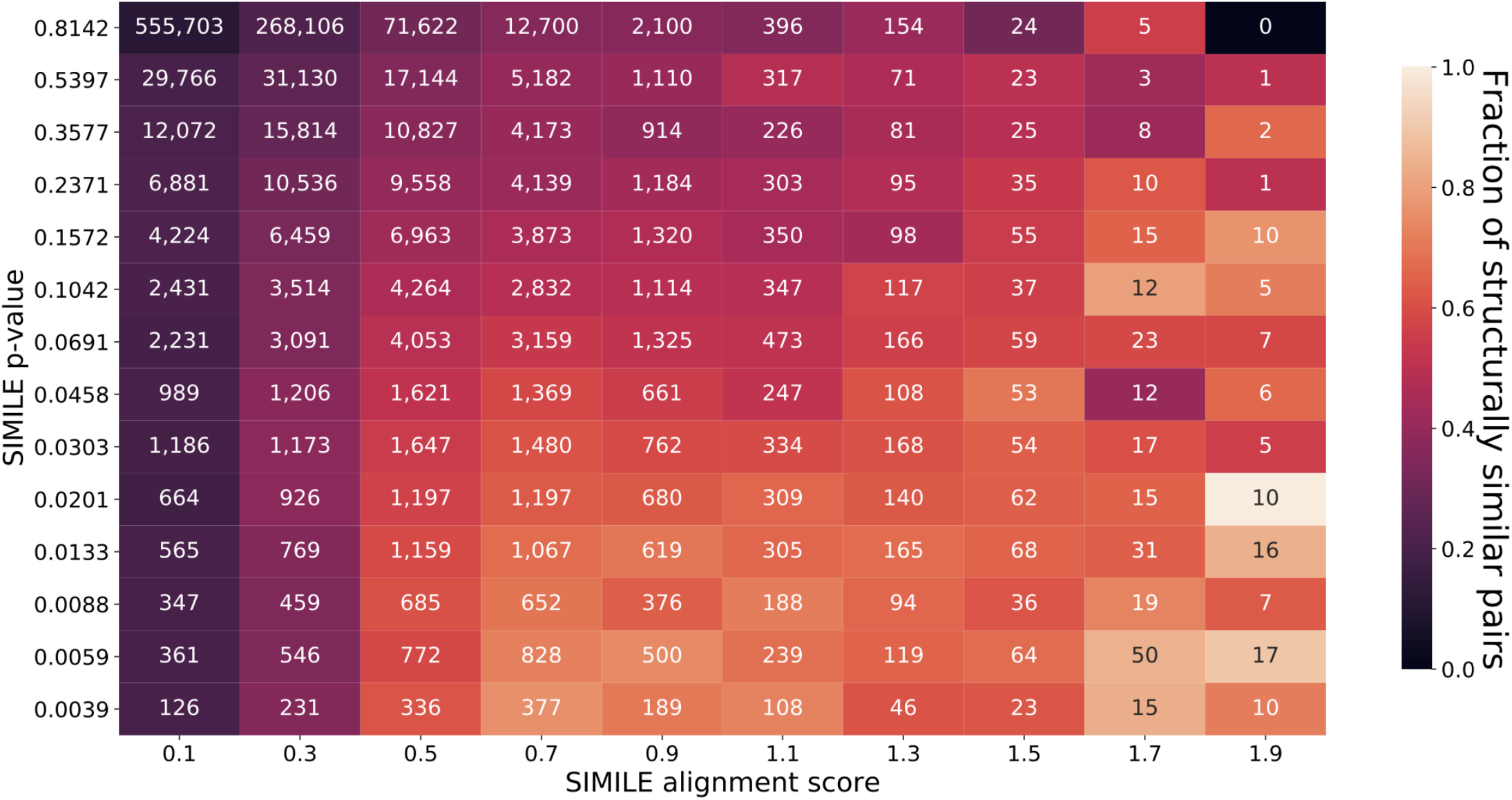
Within each window of simile alignment scores and p-values, shown is the fraction of structurally related compounds (colorscape) and the number of pairs of spectra with MCS≥0.4 (text). For higher alignment scores and lower p-values an increasing trend for both fraction of structurally related compounds. This specificity comes at the loss of sensitivity which drops dramatically with the use of strict filtering.

### Chemical class dependencies

Comparison of the number of matches enabled by SIMILE vs. modified cosine can be seen in Fig 4 as a function of compound class. Unlike SIMILE, modified cosine requires filtering identifications by matching ions. Increasing the number of matching ions from 5 to 10 greatly increases the number of structurally similar hits when using modified cosine (Fig. 4) For SIMILE, filtering by the number of matching ions aids the alignment score to a lesser extent. Cutoffs under all conditions are optimized to maximize the number of returned structurally similar compounds (MCS≥0.4) under the restriction that at least 75% of what is returned is structurally similar. When constrained by matching peaks greater than or equal to 10 for modified cosine and 10 or more aligned ions for SIMILE, modified cosine returned 39,000 structurally similar pairs of compounds while SIMILE returned 28,000. With a constraint of five or more matching ions, modified cosine returned 3,300 structurally similar pairs of compounds while SIMILE returned 22,000. Finally, without a constraint for the number of matching ions, modified cosine returned 116 structurally similar pairs of compounds while SIMILE returned 20,000.

**Fig. 4.**
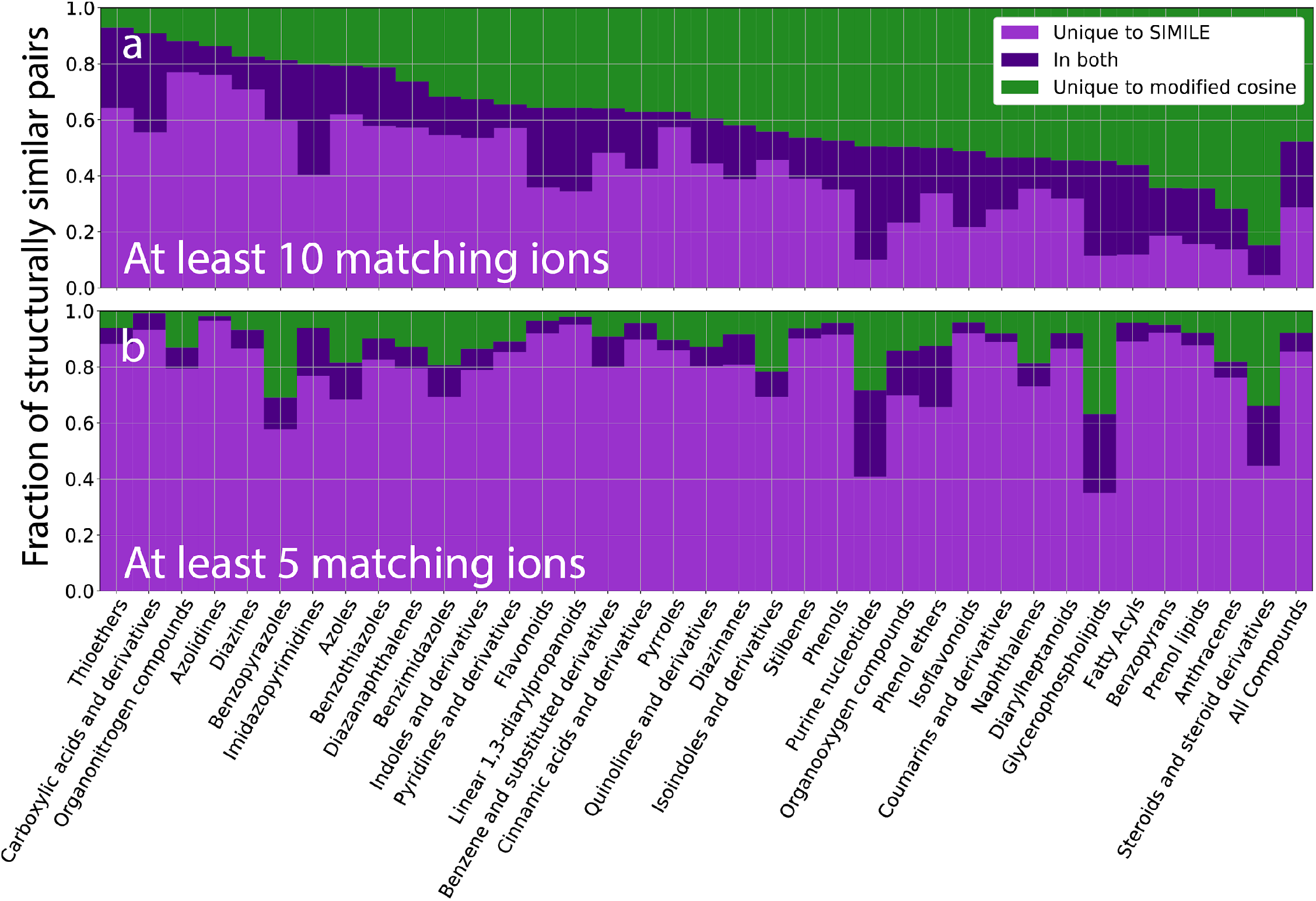
The SIMILE algorithm can identify related compounds that would be missed by existing approaches. For each class of compounds the intersection and differences of structurally similar hits (MCS>0.4) for both SIMILE and modified cosine are shown. For both modified cosine and SIMILE, scorin-parameters were used that give on average 75% of the hits with MCS of 0.4 or greater. The compound classes are provided by Classyfire. Only classes with at least 200,000 comparisons were considered where one or both compounds associated by these approaches must be in a particular compound class. a) minimum of 10 matching peaks for modified cosine or SIMILE; b) minimum of 5 matching peaks for modified cosine or SIMILE.

For the majority of compound classes, the two methods typically produced different pairs of compounds and had small overlap. In general, the number of positive identifications achievable by modified cosine is determined by the number of matching peaks. On average across compound classes, with 10 matching peaks required 40% were unique to SIMILE; 40% were unique to modified cosine; and 20% were common to both (Fig 4a). With 5 matching peaks required, 78% were unique to SIMILE; 13% were unique to modified cosine; and 9% were common to both (Fig 4b). In the case where matching peaks is not used as a filter, the structurally similar pairs of compounds returned were almost exclusively from SIMILE (greater than 99%).

### Alignment analysis

Since this is the first application of a pair-specific substitution matrix for fragmentation spectra, we sought to better understand examples where SIMILE performed remarkably well and where SIMILE performed poorly. To look closely at cases that illustrate certain strengths and weaknesses of the current implementation of SIMILE, two cases were selected and shown in Fig 5.

**Fig. 5.**
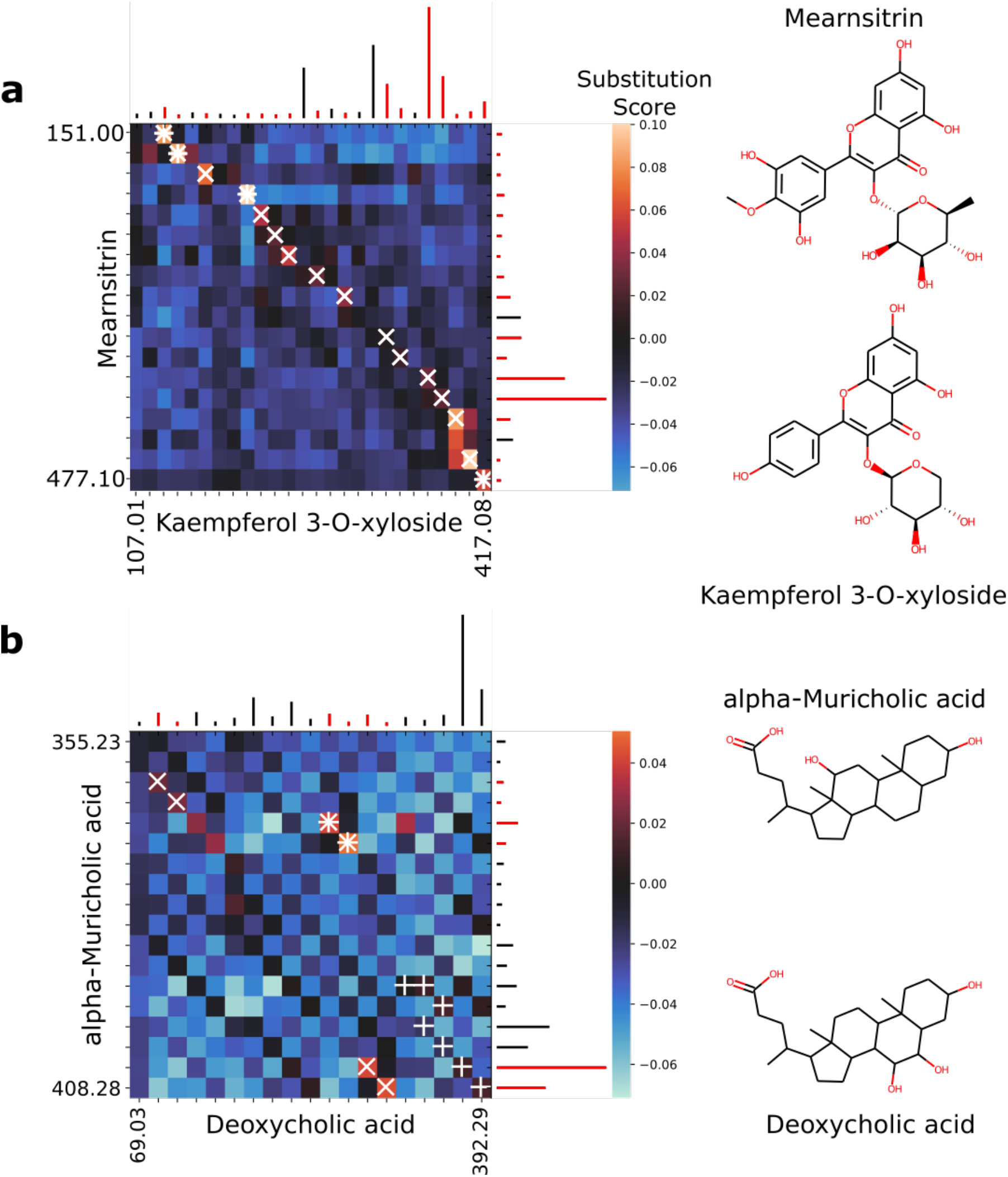
Two cases were selected to show detailed aspects of SIMILE alignment and scoring. Fragment ions aligned by both SIMILE and modified cosine are denoted by **\***; only aligned by SIMILE as **X**; and only aligned by modified cosine as **+**. Fragment ions identified by SIMILE are shown in red alongside the substitution matrix. The *m/z* differences in SIMILE alignments can correspond to compound structural differences unobservable by only considering differences in precursor ion neutral-losses. Alternatively, SIMILE alignments may underperform modified cosine when the optimal alignment relies on precursor ion neutral-losses. **a)** For a case where SIMILE dramatically outperforms modified cosine, none of *m/z* differences correspond to precursor ion neutral-losses and so the difference in the number of matching ions is extreme (4 vs 16). Additionally, 11 of aligned fragment ions have an *m/z* difference of 30.983 which likely corresponds to a chemical difference of [+O2 −H]. **b)** For a case where modified cosine dramatically outperforms SIMILE, a precursor ion neutral-loss alignment yields more matching fragment ions than SIMILE (7 vs 6). Additionally, SIMILE does not align the highest intensity ions together whereas modified cosine does.

Shown in Fig 5a is a case comparing spectra from two flavonoid molecules with high structural similarity (MCS of 0.89) but differing by four relatively small non-stereo-specific structural differences. In this example, there is a spectrum with 26 ions from kaempferol 3-O-xyloside and a spectrum with 18 ions from mearnsitrin. The alignment identified by SIMILE follows a nearly unbroken path using 16 out of a possible 18 fragment ions, with an alignment score of 0.84 and a p-value of 0.008. In line with the view that modified cosine can struggle to align compounds with multiple structural differences, modified cosine only aligned 4 ions and had a score near zero. This example demonstrates a case where a precursor ion neutral-loss shift between the two spectra fails to align the spectra, but traversing the pair-specific substitution matrix with a dynamic programming approach yields an excellent alignment.

Shown in Fig 5b is a case comparing the spectra from two secondary bile acids (deoxycholic acid with 19 ions and alpha-muricholic acid with 18 ions) differing by two structural modifications: the addition of a hydroxyl group and the relocation of another hydroxyl group. This pair of molecules has an MCS of 0.91. Like the example above, these spectra come from compounds with high overall structural similarity and differ by more than one structural difference. However, in this case SIMILE failed to find reliable alignment. Only 6 out of 18 ions were aligned with an alignment score of 0.22 and a p-value of 1. By comparison, modified cosine aligned 7 ions including the top two ions resulting in a score of 0.84. As can be seen in the substitution matrix, there are two potential paths. One path shifted by the precursor ion neutral-loss and one path based on *m/z* differences of the fragment ions. Likely an improvement to the SIMILE algorithm that incorporates precursor ion neutral-loss differences (like modified cosine) would aid this case and potentially many others.

## Discussion

Spectral similarity applications, like spectral networking, benefits from increasing connections between truly similar metabolites. One way of accomplishing this is by using a scoring method that is robust to arbitrary structural differences and provides statistical significance. In protein sequence alignment, this is achieved by using substitution matrices with alignment algorithms that generate scores according to known distributions^19–21^. However, substitution matrices also appear in many other contexts including text, speech, video, or even mathematically abstracted signals^22–25^. Once a substitution matrix is chosen, widely known methods can be used to calculate optimal alignments between pairs of signals^9,14,26^. To this end, we have developed a method of generating substitution matrices for fragmentation spectra to associate fragment ions by the similarity of their fragmentation paths. We found that our alignment method, SIMILE, yields different and typically more associations than modified cosine, significant alignment scores generated by SIMILE correspond to compound structural similarity, and the effectiveness depends on compound class.

As can be seen in Figure 4a, when filtered by a minimum of 10 matching ions, SIMILE performs comparably to modified cosine in detecting long-range structural similarity between compounds, but typically identifies different pairs of compounds. Additionally, the relative performance of SIMILE to modified cosine is stratified by compound class. This provides evidence that the SIMILE alignment scores and p-values are capturing aspects of how similar compounds fragment similarly. For modified cosine, the number of matching ions acts as a heuristic to approximate the significance of aligning/scoring the similarity of two spectra; in contrast with SIMILE, there is a mechanistic calculation of significance using a framework based on fragment ion substitutability. Likely there will be classes of molecules that are more appropriate for cosine based scoring than SIMILE based scoring. For example, as shown in Fig 4, the SIMILE algorithm does not perform as well for steroids and glycerophospholipids as it does for other classes of compounds. Since steroids and glycerophospholipids are very common classes of compounds in both reference libraries and in experimental observations, it makes sense to use SIMILE as an algorithm to accompany traditional scoring approaches (like modified cosine).

The underlying SIMILE distance measure for comparing fragment ions is closely related to Euclidean Commute Time Distance (ECTD), which has the property of decreasing with the number of connecting paths and increasing with the “length” of connecting paths^27^. The number of paths connecting two fragment ions increases when the number of total fragment ions increases. Likewise, the “length” of a path connecting fragment ions decreases when the frequency of m/z differences in the path increases. In other words, two fragment ions are similar if they are connected by many paths exhibiting high frequency *m/z* differences.

Saerens et al. prove the pseudo-inverse of the unnormalized graph laplacian acts as a covariance matrix with respect to ECTD^27^. We use a normalized and directed graph laplacian as the SIMILE substitution matrix to ensure substitution scores in the fragment ion substitution matrix are normalized and the asymmetry of substitution is captured. We can then score fragmentation spectra pairs by using a dynamic programming algorithm to align their fragment ions according to substitution scores^14^. This in turn mirrors how pairs of protein sequences can be scored by aligning their residues according to a substitution matrix.

For much of metabolomics research (including spectral networking), it is necessary to use intermediate spectral similarity scores. Interrelating fragmentation spectra with intermediate scores in SIMILE is aided by a significance estimate; without statistical significance, intermediate similarity scores are to some degree uninterpretable. For example, intermediate scores could imply moderate structural similarity or tentative high similarity. Interpretation of intermediate scores has a direct effect on how researchers prioritize metabolites of interest and ultimately on the outcome of their research. As such, SIMILE significance estimation assists in deriving biological insight from metabolomics data when confronted with underexplored biochemistry.

Significance of SIMILE alignments are calculated via a Monte Carlo permutation test with alignment score as the test statistic. It is important to note that significant SIMILE alignment of fragmentation spectra does not necessitate compound structure similarity. This is similar to how protein sequences can have significant alignment but yield different tertiary structures or functions. Likely a significant SIMILE alignment indicates that two molecules fragment similarly, not that they necessarily have similar structures. Nevertheless, Fig 3 and Fig 4 show significant fragmentation spectra alignments often correspond to moderate to high structural similarity.

While SIMILE is a very promising new approach for scoring pairwise spectral alignments, since this is the first application of scoring fragmentation spectra using a substitution matrix, close attention must be applied. We recommend complementing SIMILE with the use of cosine based scoring for compound identification. There is a massive amount of literature on cosine based scoring, is well established, and has been in use for many decades. Since we find that SIMILE often works for cases where cosine based scoring fails, using both approaches will provide more identifications. In addition, SIMILE provides a p-value and an alignment between fragments. From the m/z differences in the alignment, one can glean structural clues regarding the differences between the molecules. For example, the predominant *m/z* difference in Fig 5a of 30.983 likely corresponds to [+O2 −H]. This is not surprising given one of the chemical differences between the two structures is the addition of two oxygens. We see this methodology being useful for elucidation of novel natural products by using the fragmentation spectra of known members of the same chemical class as “scaffolds.” In addition, as shown in Fig 5b, there are improvements to the SIMILE algorithm that can likely boost its reliability and interpretability.

## Conclusion

Here we describe SIMILE, a metabolomics tool immediately useful to complement existing widely used approaches with the potential to open up a completely new area for research in fragment ion substitution matrices with significance estimation. This approach provides a new scoring and significance framework for discovering relationships between molecules that would have been dismissed with existing approaches. As such, using SIMILE as an algorithm to accompany traditional scoring approaches (like modified cosine) should lead to increased discovery in multiple fields including compound and pathway discovery, and other useful applications of spectral networking.

## Acknowledgements

This work was supported by the U.S. Department of Energy Office of Science by, ENIGMA-Ecosystems and Networks Integrated with Genes and Molecular Assemblies (http://enigma.lbl.gov), a Science Focus Area Program at Lawrence Berkeley National Laboratory; the U.S. Department of Energy Joint Genome Institute (JGI) a DOE Office of Science User Facility; the Machine Learning for Chemistry Laboratory Directed Research and Development Program; and the National Energy Research Scientific Computing Center (NERSC) a DOE Office of Science User Facility all under Contract No. DE-AC02-05CH11231 with the U.S. Department of Energy. The U.S. Government retains, and the publisher, by accepting the article for publication, acknowledges, that the U.S. Government retains a non-exclusive, paid-up, irrevocable, world-wide license to publish or reproduce the published form of this manuscript, or allow others to do so, for U.S. Government purposes.

